# Lipid Droplets in Brown Adipose Tissue Are Dispensable for Cold-Induced Thermogenesis

**DOI:** 10.1101/2020.03.30.016451

**Authors:** Chandramohan Chitraju, Alexander Fischer, Robert V. Farese, Tobias C. Walther

**Affiliations:** Department of Molecular Metabolism, Harvard T.H. Chan School of Public Health, Boston, MA 02115, USA; Department of Cell Biology, Harvard Medical School, Boston, MA 02115, USA; Broad Institute of Harvard and MIT, Cambridge, MA 02142, USA; Howard Hughes Medical Institute, Boston, MA 02115, USA

## Abstract

Brown adipocytes store metabolic energy as triglycerides (TG) in multilocular lipid droplets (LDs). Fatty acids released from brown adipocyte LDs by lipolysis are thought to activate and fuel UCP1-mediated thermogenesis. Here we test this hypothesis by preventing fatty acid storage in murine brown adipocytes through brown adipose tissue (BAT)-specific deletions of the TG synthesis enzymes, DGAT1 and DGAT2 (BA-DGAT KO). Despite the absence of LDs, BA-DGAT KO mice had functional BAT and maintained euthermia during acute or chronic cold exposure. As apparent adaptations to the lack of TG, brown adipocytes of BA-DGAT KO mice appear to utilize circulating glucose and fatty acids, as well as stored glycogen to fuel thermogenesis. Moreover, BA-DGAT KO mice were resistant to diet-induced glucose intolerance, likely due to increased glucose disposal by BAT. Thus, surprisingly, TGs in BAT are dispensable for its function, in part through adaptations to utilize other fuel sources.

## INTRODUCTION

Homeotherms maintain constant body temperature despite changes in ambient temperature. Mammals maintain core body temperature by adaptive thermogenesis, including shivering and non–shivering thermogenesis (Cannon and Nedergaard, 2004; Lowell and Spiegelman, 2000). In non-shivering thermogenesis, brown and beige fat dissipate chemical energy as heat by uncoupling respiration from ATP synthesis (Smith and Roberts, 1964; Smith et al., 1966; Wu et al., 2012) as well as by other futile enzymatic cycles (Chouchani et al., 2019; Kazak et al., 2015).

Non-shivering thermogenesis is fuelled by different sources, including glucose, succinate (Mills et al., 2018), branched-chain amino acids (Yoneshiro et al., 2019), and fatty acids (FAs) (Cannon and Nedergaard, 2004; Townsend and Tseng, 2014). In rodents, glucose is a major fuel, with ∼20% of circulating glucose being consumed by BAT under basal conditions (Hankir and Klingenspor, 2018). FAs, derived from triglycerides (TGs) stored in lipid droplets (LDs) of brown adipocytes, from white adipose tissue (WAT) lipolysis (Schreiber et al., 2017), or from TG-rich lipoproteins in the circulation (Bartelt et al., 2011), are other major fuels. Cold exposure of mice triggers activation of the sympathetic nervous system, which in turn activates adipose TG lipase (ATGL)-mediated lipolysis of TGs stored in LDs of brown and white adipocytes to liberate FAs (Schreiber et al., 2017; Zechner et al., 2012). The requirement for FAs stored in TGs of brown adipocytes for thermogenesis has not been determined. Recent studies showed that lipolysis catalyzed by ATGL in brown adipocytes is not required for mice to maintain body temperature during cold exposure (Schreiber et al., 2017; Shin et al., 2017). This suggests that FAs stored in brown adipocytes are not strictly required. However, since hormone-sensitive lipase (HSL) can also catalyze TG hydrolysis, alternative mechanisms of lipolysis may have compensated for loss of ATGL in these studies.

Here we sought to determine whether TG storage in LDs of brown adipocytes is required for thermogenesis and euthermia. To address this, we generated mice lacking TGs in BAT by deleting both acyl CoA:diacylglycerol acyltransferase (DGAT) enzymes, DGAT1 and DGAT2 (Cases et al., 1998; Cases et al., 2001; Yen et al., 2008), specifically in BAT (BA-DGAT KO). We then studied the physiology of these mice in response to thermogenic challenges.

## RESULTS

### Triglyceride Stores and Lipid Droplets Are Absent in Brown Adipocytes of Mice Lacking Both DGAT1 and DGAT2

To generate BAT-specific *Dgat1* and *Dgat2* double knockout (BA-DGAT KO) mice, we first generated *Dgat1* and *Dgat2* double-floxed mice (D1D2 flox) by crossing *Dgat1*^*flox/flox*^ mice (Shih et al., 2009) with *Dgat2*^*flox/flox*^ mice (Chitraju et al., 2019). We then crossed D1D2 flox mice with transgenic mice expressing Cre recombinase under control of the murine *Ucp1* promoter (Kong et al., 2014).

BA-DGAT KO mice were healthy and yielded offspring with the predicted Mendelian ratio of genotypes. *Dgat1* and *Dgat2* mRNA levels were decreased by ∼95% and ∼85%, respectively, in interscapular brown adipose tissue (BAT) of BA-DGAT KO mice but were unaltered in inguinal white adipose tissue (iWAT) (Figure 1A). DGAT activity was decreased by ∼95% in BAT of BA-DGAT KO (Figure 1B). Dual-energy X-ray absorptiometry (DEXA) analysis showed that BA-DGAT KO mice and control D1D2 flox mice had similar fat and lean masses overall (Figure 1C), which was also reflected in experiments with nuclear magnetic resonance imaging (Figure S1A). Weights of gonadal WAT depots (Figure S1B) were also similar in control D1D2 flox and BA-DGAT KO mice. However, BAT from BA-DGAT KO mice appeared darker brown than BAT from control mice (Figure 1D) and was more dense, sinking in a liquid fixative (Figure 1E). Analysis of the lipids in the BAT-DGAT KO confirmed that TG stores in BAT were reduced by ∼95% (Figure 1F). Histological examination revealed that LDs were absent in nearly all cells of BAT depots specifically in BA-DGAT KO mice (Figure 1G). A few cells of unknown identity, which presumably do not express UCP1-Cre, in BAT of BA-DGAT KO mice had LDs (Figure 1G). These cells may account for the little TG present in BAT of BA-DGAT KO mice. Transmission electron microscopy (TEM) analyses verified that brown adipocytes of BA-DGAT KO mice lacked LDs but had abundant mitochondria (Figure 1G). Plasma levels of TGs were ∼15% reduced in *ad libitum* fed BA-DGAT KO mice (41±8 mg/dl vs. 31±4 mg/dl, respectively, p=0.04) and glucose levels were slightly lower (158 ± 9 mg/dl vs. 146 ± 8 mg/ml, respectively, p=0.01) than in control mice (Figures S1C and S1D). Levels of free FAs in plasma were not different (Figure S1C).

**Figure 1.**
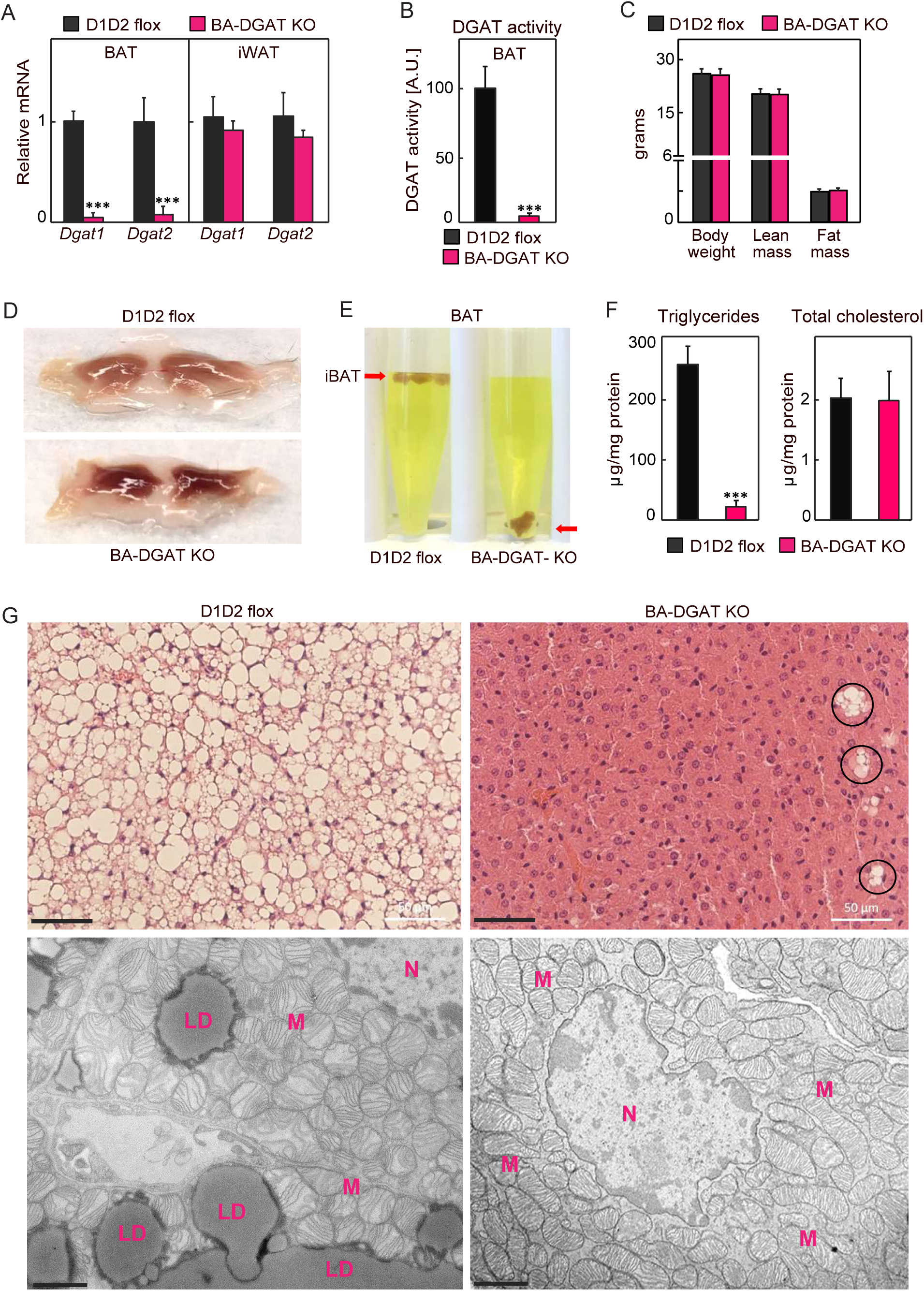
Triglycerides and Lipid Droplets (LDs) Are Absent in BAT of BA-DGAT KO Mice. Brown adipose tissue specific *Dgat1* and *Dgat2* double knockout mice (BA-DGAT KO) were generated by crossing D1D2 flox mice with the mice expressing Cre-recombinase under control of the *Ucp1* promoter. (A) mRNA levels of *Dgat1* and *Dgat2* in BAT and iWAT (n=6). (B) DGAT activity in BAT lysates (n=4 mice). (C) Lean mass and fat mass analysis of 10 weeks old chow-diet fed mice (n=8). (D) Gross appearance of iBAT. (E) BAT of BA-DGAT DKO mice sinks in a liquid fixative. (F) Triglycerides and total cholesterol content in BAT (n=5 mice). (G) H&E stained sections and transmission electron microscopy (TEM) images of iBAT. Scale bars, 50 µm (H&E), 2 µm (TEM). LD, lipid droplet; M, mitochondria; N, nucleaus. Data are presented as mean ± SD. ***p<0.001 by t-test.

### BA-DGAT KO Mice Maintain Euthermia during Acute or Chronic Cold Exposure

To determine the contributions of TG stores in BAT to thermogenesis, we exposed BA-DGAT KO mice to cold temperature. In response to acute cold exposure (4°C) for 6h with *ad libitum* feeding, or for 5 hours with fasting, BA-DGAT KO mice maintained their core body temperature (Figure 2A). To further challenge thermogenic demands, we exposed mice to cold temperature for 1 week with *ad libitum* feeding. Again, we found no differences in core body temperatures between BA-DGAT KO and control mice (Figure 2A). These results indicate that TG stores of BAT are dispensable for cold-induced thermogenesis in mice.

**Figure 2.**
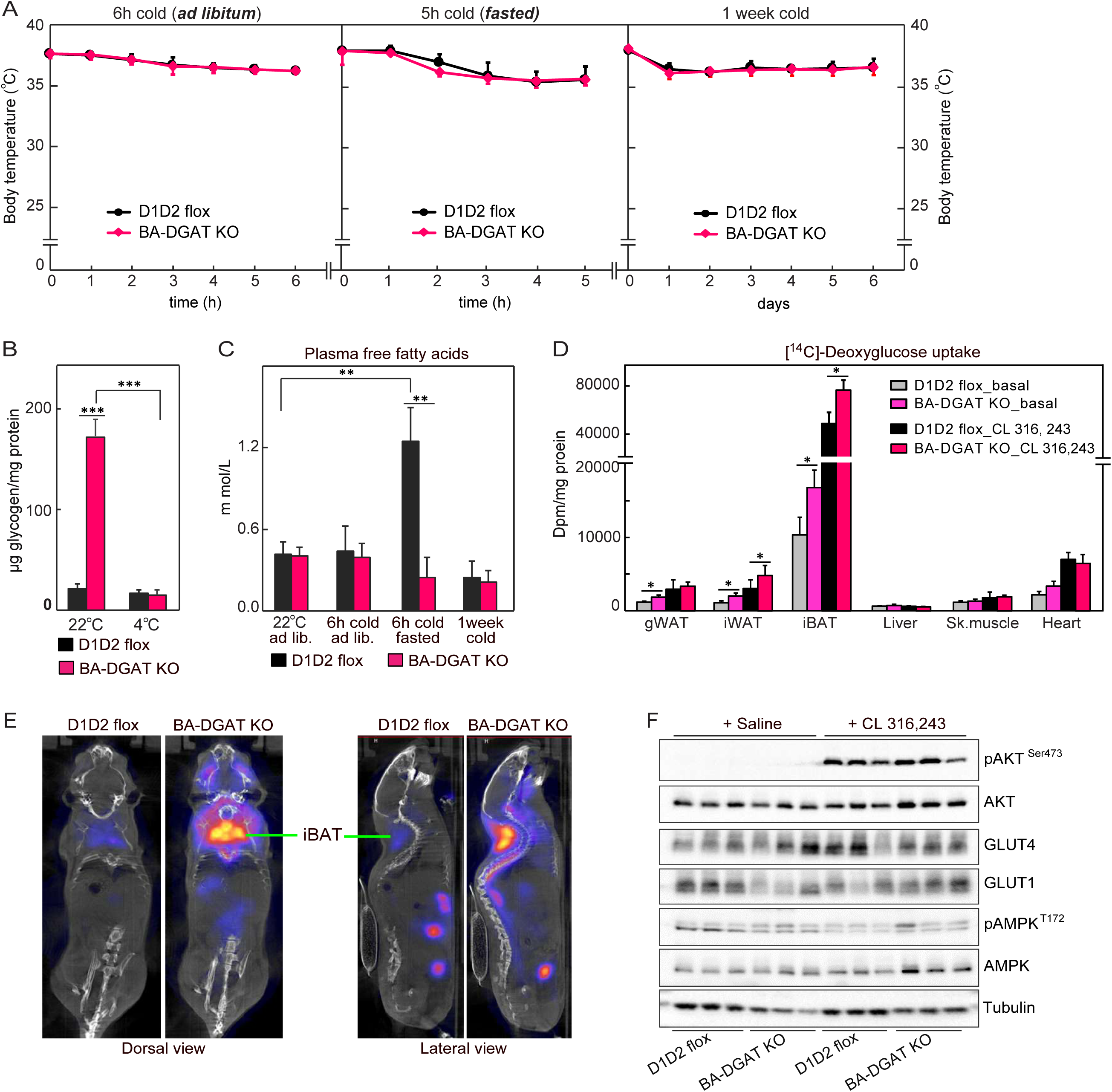
BA-DGAT KO Mice Maintain Euthermia during Acute or Chronic Cold Exposure. (A) Mice were exposed to cold acutely (in *ad libitum* fed or in fasted state) or chronically for a week (n=8 mice). (B) Glycogen levels in brown fat (n=3 mice). (C) Plasma free fatty acids (n=7 mice). (D) [^14^C]-Deoxyglucose uptake by tissues *in vivo* (n=3 mice). (E) ^18^-FDG-PET/CT scans of CL 316,243-injected mice. (F) Western blot analysis of insulin signaling in BAT of basal or CL 316,243-injected mice (n=3 mice). Data are presented as mean ± SD. *p<0.05, **p<0.01, ***p<0.001.

We hypothesized that brown adipocytes of BA-DGAT KO mice utilize other sources of fuel to enable thermogenesis. Tissue glycogen levels were ∼5-fold higher in BAT of BA-DGAT KO mice housed at room temperature, and the excess glycogen was no longer apparent after 6 hours of cold exposure (Figure 2B), suggesting that glycogen is a compensatory fuel. In agreement with this, we found moderate increases in the mRNA levels of glycogen synthase and glycogenin in BAT of BA-DGAT KO mice (see Figure 3A).

**Figure 3.**
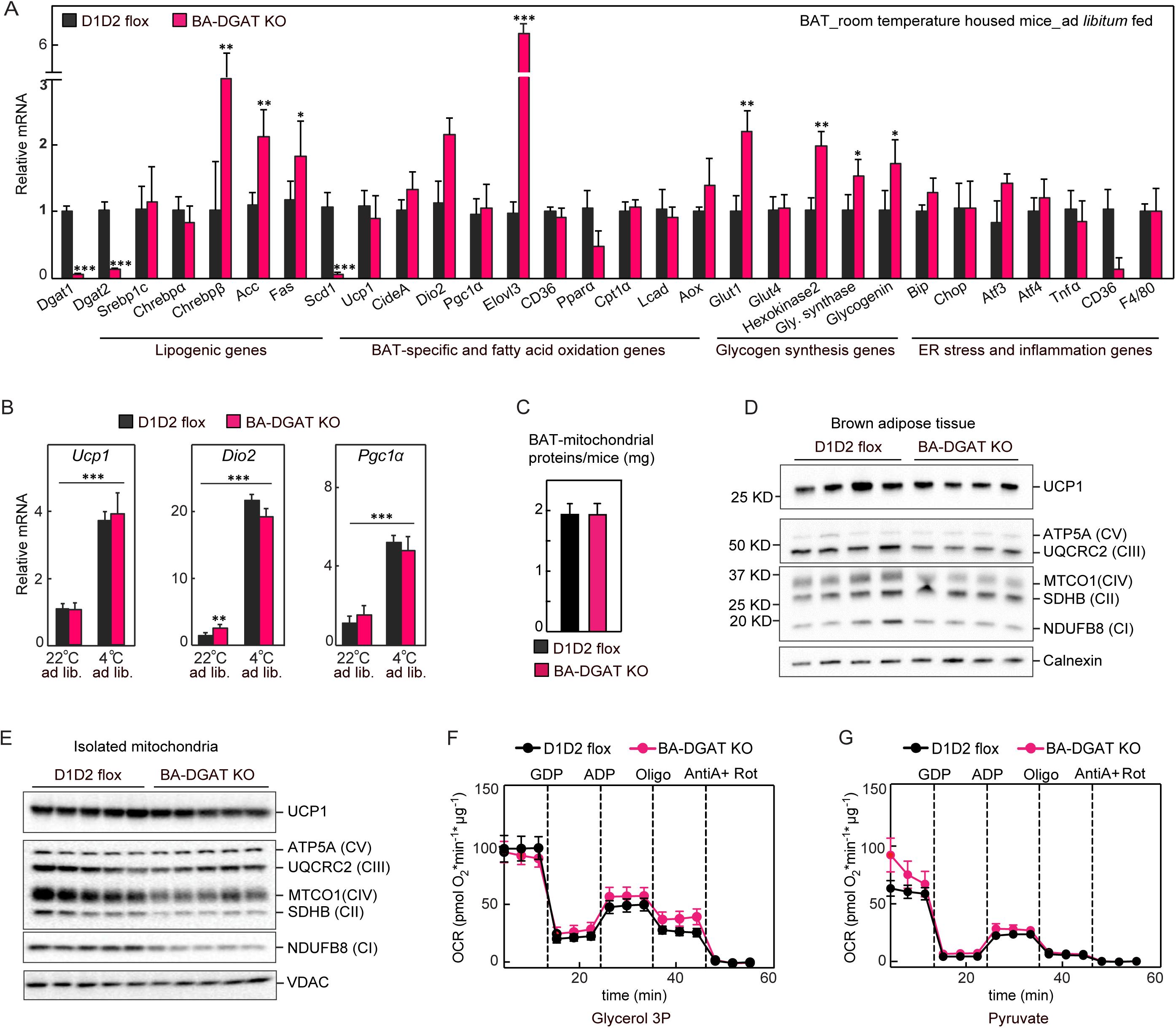
BAT Lacking TG Stores Exhibit Adaptations to Maintain Thermogenesis. (A) mRNA levels in BAT of mice housed at room temperature (n=6 mice). (B) mRNA levels in BAT of *ad libitum* fed or cold-exposed (fasted) mice (n=6 mice). (C) Total mitochondrial protein content in isolated mitochondria (n=8 mice). (D) Western blot analysis of mitochondrial proteins from BAT tissue lysates (n=4 mice). (E) Western blot analysis of mitochondrial proteins from isolated mitochondria (n=4 mice). (F and G) oxygen consumption rates (OCR) of isolated mitochondria measured using glycerol-3-phosphate or pyruvate. The difference between initial respiration and respiration after GDP addition was considered UCP1-activity. The difference between respiration after ADP addition and after oligomycin addition was considered to be ATP synthase activity. Data are presented as mean ± SD. *p<0.05, **p<0.01, ***p<0.001.

BAT can also utilize circulating glucose and FAs to fuel cold-induced thermogenesis (Cannon and Nedergaard, 2004; Townsend and Tseng, 2014). For example, FAs released from WAT can fuel BAT-mediated thermogenesis in the fasted state (Schreiber et al., 2017). Consistent with this, FA levels were increased two-to threefold in fasted control mice exposed to cold (Figure 2C). This increase was not found in fasted BA-DGAT KO mice, suggesting that circulating FAs were being utilized for thermogenesis in BAT. Consistent with this hypothesis, levels of [^14^C]-oleic acid that were taken up and accumulated in BAT were reduced in BA-DGAT KO mice, suggesting increased fat oxidation (Figure S3C).

We also determined if glucose uptake by BAT was altered in BA-DGAT KO mice. For these experiments, we treated mice with a β-3-adrenoceptor agonist (CL 316,243), which activates thermogenesis and increases glucose uptake (Cannon and Nedergaard, 2004). Indeed, [^14^C]-deoxyglucose uptake by BAT of BA-DGAT KO mice was ∼30% greater in the basal state or after treatment with CL 316,243 than in controls (Figure 2D). Similarly, ^18^-fluoro-deoxyglucose positron emission tomography-computed tomography scanning (^18^-FDG-PET/CT) of mice after CL 316,243 injection showed that BAT of BA-DGAT KO mice took up more glucose than BAT of control mice (4.2E+08 Bqml/mm^3^ vs. 2.2E+08 Bqml/mm^3^, respectively, (Figure 2E)).

We investigated the mechanisms underlying the increased glucose uptake in BA-DGAT KO mice (Figures 2F and 3A-B). β-Adrenergic signaling in brown adipocytes activates AKT (Heine et al., 2018; Sanchez-Gurmaches et al., 2018), and we found that CL 316,243 stimulated phosphorylation of AKT (pAKT^Ser473^) similarly in BAT of BA-DGAT KO and control mice (Figures 2F). Consistent with this finding, phosphorylation of AKT (pAKT^Ser473^) was also similar in BAT of BA-DGAT KO and control mice after insulin treatment (Figure S3B). The mRNA expression of the glucose transporter, *Glut1*, was increased twofold in BAT of BA-DGAT KO mice, but GLUT1 protein levels were similar in BA-DGAT KO and control mice. *Glut4* mRNA and protein levels were not changed. These findings suggest no gross alterations of glucose transporter expression, but they do not exclude the possibility that more transporters may be localized to the plasma membranes of brown adipocytes in BA-DGAT KO mice. AMPK and thyroid hormone also regulate BAT glucose uptake by an insulin-independent mechanism (de Jesus et al., 2001; Teperino et al., 2012). We found no differences in pAMPK^T172^ levels in BAT of BA-DGAT KO and control mice (Figure 2F). Expression levels of type II deiodinase, *Dio2*, which coverts thyroxine (T4) to bioactive 3,3’,5-triiodothyronine (T3), were increased in BAT of room temperature-housed BA-DGAT KO mice (Figures 3A, and 3B), but were not increased in BAT of cold-exposed BA-DGAT KO mice.

We also performed indirect calorimetry studies after activation of β-3-adrenoceptor by CL 316,243 treatment. Basal rates of oxygen consumption, CO_2_ production, and respiratory exchange ratio (RER) were normal in BA-DGAT KO mice (Figure S2). However, compared with control mice, BA-DGAT KO mice exhibited a reduced increase in oxygen consumption and energy expenditure after CL 316,243 injection, suggesting that BAT with TG stores has a higher respiratory capacity than BAT without lipid stores. Consistent with this interpretation, the RER of BA-DGAT KO mice was slightly greater than controls after CL 316,243 treatment, suggesting BA-DGAT KO mice utilize less fat or more glucose in response to this challenge (Figure S2).

### BAT Lacking TG Stores Exhibits Adaptations to Maintain Thermogenesis

Our results indicate that BAT lacking TG stores can compensate to fuel cold-induced thermogenesis in part by utilizing alternative fuels, such as tissue glycogen and circulating glucose and FAs. We reasoned that adaptations in BAT function may also occur in BA-DGAT KO mice. To examine this possibility, we analyzed the expression of thermogenic genes and proteins. At baseline or with cold-exposure, BAT of BA-DGAT KO mice exhibited gene expression changes, consistent with increased *de novo* FA synthesis (*e.g*., *Chrebpβ, Acc, Fas*) and elongation (*Elovl3*), although the expression of the Δ9 desaturase *Scd1* was markedly decreased (Figure 3A). Upon cold exposure, activation of expression of BAT-specific and thermogenic genes, such as *Ucp1, Pgc1α, CideA, and Dio2*, was similar in BAT of BA-DGAT KO and control mice (Figure 3B), as were the mRNA expression levels of genes involved in FA oxidation (Figures 3A, 3B and S3A). Storing TG in LDs protects white adipocytes from lipid-ER stress (Chitraju et al., 2017). Expression of genes of the ER stress response were not increased in BA-DGAT KO mice under these conditions (Figures 3A and S3A), consistent with the notion of FAs being oxidized and not accumulating in membranes. These findings suggest adaptive changes in BAT of BA-DGAT KO mice that promote the synthesis of long-chain FAs, possibly to drive UCP1 activation and as a source for oxidative fuel.

Mitochondrial morphology, as assessed by EM, appeared to be grossly normal in brown adipocytes of BA-DGAT KO mice (Figure 1G), and mitochondrial cross-sectional area was similar (Figure S1F). Total potein levels were similar in mitochondria isolated from BAT of BA-DGAT KO and control mice (Figure 3C). UCP1 protein levels in whole BAT tissue lysates and in isolated mitochondria of BA-DGAT KO mice were normal (Figures 3D and 3E). Western blot analysis of respiratory complex proteins showed moderate reductions in NDUFB8 (complex I) and MTCO1 (complex IV) proteins in whole BAT tissue lysates and isolated mitochondria of BA-DGAT KO mice (Figures 3D and 3E). Complex II levels were also mildly reduced in isolated mitochondria of BA-DGAT KO mice (Figure 3E). However, we found similar levels of UQCRC2 (complex III) and ATPA (complex V) (Figures 3D and 3E).

To test whether the changes in mitochondrial proteins affect mitochondrial respiration, we isolated mitochondria from BAT and measured oxygen consumption rates (OCR) in response to different substrates. The basal OCR was similar in mitochondria isolated from BA-DGAT KO and control mice with glycerol-3-phosphate or pyruvate as substrates (Figures 3F and 3G). Respiration with maximal ATP-synthase activity (ADP addition) was slightly reduced in mitochondria of BA-DGAT KO mice. UCP1-dependent respiration (GDP-inhibitable) was similar in mitochondria of BA-DGAT KO and controls with glycerol-3-phosphate substrate or with pyruvate (Figures 3F and 3G).

These results suggest that while mitochondria of BAT of BA-DGAT KO mice exhibit some changes in levels of OXPHOS complex proteins, they appear to have adapted to maintain relatively normal mitochondrial respiration..

### BA-DGAT KO Mice Are Resistant to High-Fat-Diet-Induced Glucose Intolerance

Given increased glucose uptake into BAT of BA-DGAT KO mice, we hypothesized that BA-DGAT KO mice may have improved glucose tolerance under conditions that promote insulin resistance. To test this hypothesis, we fed mice a high-fat diet (HFD) for 12 weeks. At the end of the study period, BAT from BA-DGAT KO mice exhibited a phenotype similar to BAT of chow-fed BA-DGAT KO mice, including a darker color (Figure S4A), higher density in a liquid fixative (Figure S4B), and near absence of LDs (Figure 4A). Also, a few cells of unknown identity in BAT of BA-DGAT KO mice had LDs (Figure 4A). Glycogen levels were ∼10 fold higher in BAT of HFD-fed BA-DGAT KO mice (Figure 4B). Glycogen granules were visible in electron microscopy images of BAT from HFD-fed BA-DGAT KO mice (Figures 4C and S4C). Since previous studies showed that DGAT-catalyzed TG synthesis is important to protect cells from high intracellular FA levels (Chitraju et al., 2017; Koliwad et al., 2010; Listenberger et al., 2003; Liu et al., 2014), we measured expression ER stress and inflammatory marker genes but found no increases (Figure 4G).

**Figure 4.**
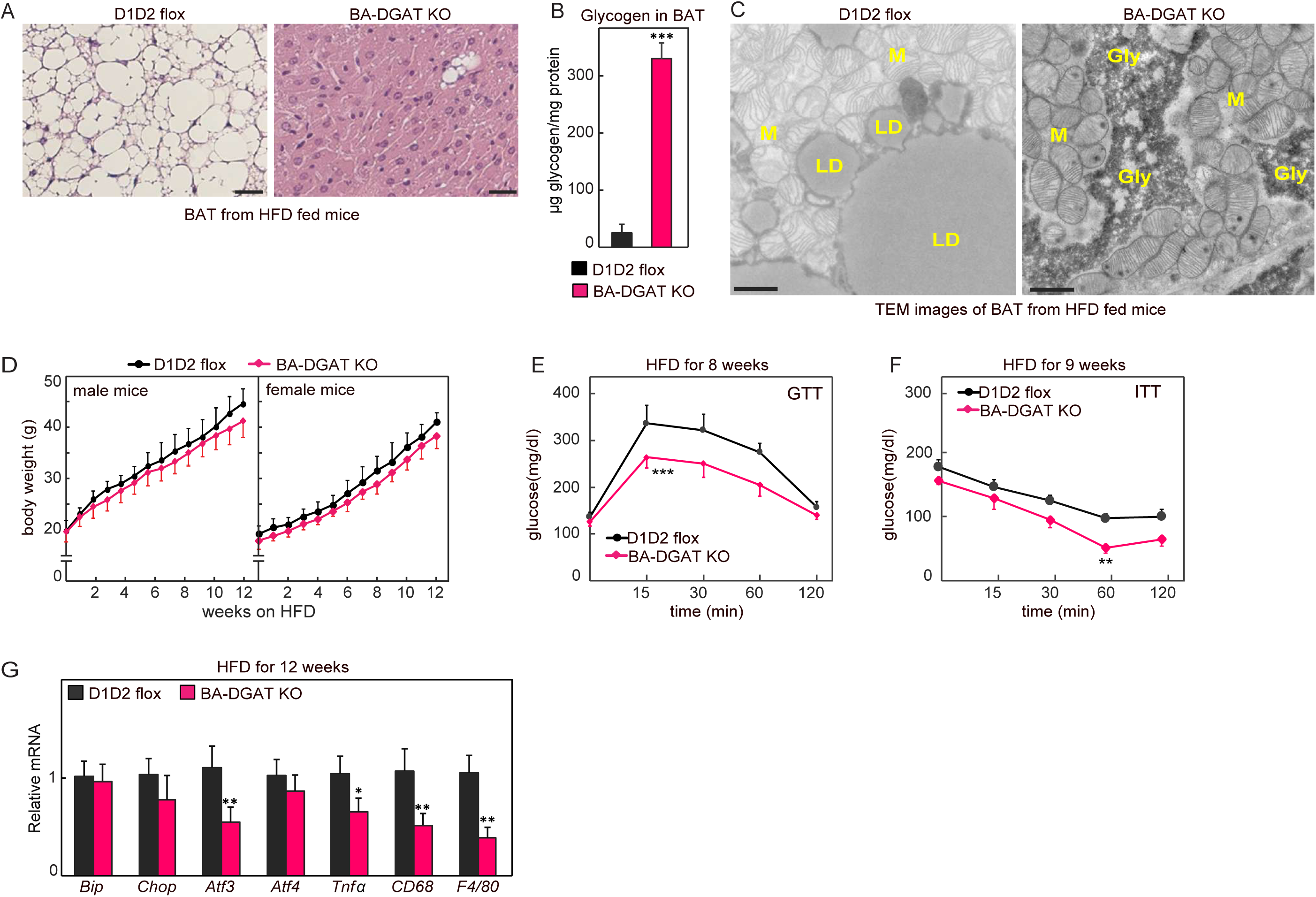
BA-DGAT KO Mice Are Resistant to High-fat-diet (HFD)-Induced Glucose Intolerance. (A) H&E-stained sections of iBAT from 12 weeks HFD-fed mice, scale bars, 25 μm. (B) Glycogen levels in brown fat of HFD fed mice (n=3 mice). (C) TEM images of iBAT from HFD-fed mice, scale bars, 2 µm. LD, lipid droplet; M, mitochondria; Gly, glycogen. (D) Body weights of mice fed on Western type HFD (n=15 mice). (E and F) Glucose tolerance test and insulin tolerance test performed on HFD-fed mice (n=10). (G) mRNA levels in BAT of HFD-fed mice (n=6 mice). Data are presented as mean ± SD. **p<0.05; ***p<0.01 by two-way ANOVA (for E and F). *p<0.05, **p<0.01 by t-test (for G).

On the level of the whole animal, both control and BA-DGAT KO mice gained similar amounts of weight on the HFD (Figure 4D). However, BA-DGAT mice had enhanced glucose and insulin tolerances (Figures 4E and 4F), suggesting their increased glucose uptake into BAT protects them from the impaired glucose metabolism normally found with this diet.

## DISCUSSION

By genetically deleting both DGAT enzymes in brown adipocytes, we show here that TG storage in LDs of murine brown adipocytes is dispensable for thermogenesis in response to cold. In the absence of TG stores in brown adipocytes, other substrates, such as circulating glucose and FAs, as well as brown adipocyte glycogen, appear to be sufficient to compensate to supply fuel to drive respiration and thermogenesis. Analysis of mitochondria of brown adipocytes of BA-DGAT KO mice showed that they were mostly functioning normally. Most of the changes in brown adipocytes of BA-DGAT KO mice were in gene expression and suggest adaptations to generate long-chain saturated FAs, presumably for fuel or to drive uncoupling (Bertholet and Kirichok, 2019; Fedorenko et al., 2012). How such changes are triggered (i.e., through the SNS or other mechanisms) is unknown.

Our findings agree with those showing that ATGL-mediated lipolysis of stored TG in brown adipocytes is dispensible for cold-induced thermogenesis (Schreiber et al., 2017). A conclusion from these studies was that TG storage in brown adipocytes was likely not required for thermogenesis, provided that FAs could be derived from WAT stores. This may also be the case in our BA-DGAT KO model where TGs are still present in WAT. The two models differ, however, as the BAT of BAT-ATGL knockout mice still had TG stores that may be lipolyzed by HSL (Schweiger et al., 2006). Our model, in contrast, completely depletes TG stores in brown adipocytes and therefore definitively answers whether TG stores are required functionally.

The gene expression changes in BAT of BA-DGAT KO mice suggest that one response to a lack of TG stores in this tissue is to activate *de novo* lipogenesis. This pathway is largely regulated by SREBP and ChREBP transcription factors (Eberle et al., 2004; Horton et al., 2002; Ishii et al., 2004). The gene expression changes, with increased expression of *ChREBPβ* (a ChREBPα target), but not *Srebp1c* itself, suggest activation of the ChREBP pathway via increased glucose uptake. Indeed, these findings are consistent with glucose-mediated activation of MLX family of transcription factors (Kawaguchi et al., 2001; Ma et al., 2005), including ChREBP, and target genes that promote *de novo* lipogenesis in BAT (Mottillo et al., 2014; Sanchez-Gurmaches et al., 2018; Trayhurn, 1979; Yu et al., 2002). This finding is also consistent with our recent report of increased expression of ChREBP/MLX-target genes in cells lacking LDs through DGAT inhbition (Mejhert et al., 2020). Moreover, they agree with previous studies that suggest a feedback mechanism correlating reduced DGAT activity with inhibiton of the SREBP1-mediated lipogenic pathway (Chitraju et al., 2017; Choi et al., 2007; Monetti et al., 2007).

Finally, our studies highlight the use of glucose as a fuel for thermogenesis in rodents. We found dramatic changes in glucose upake in BAT of BA-DGAT KO mice, and these mice were protected from glucose intolerance when challenged with a high-fat diet. These findings are consistent with other studies that illustrate that glucose disposal in brown adipocytes can protect against glucose intolerance peripherally (Chondronikola et al., 2014; Lee et al., 2014; Stanford et al., 2013). These findings suggest that selective depletion of TG storage in BAT (or by extrapolation in beige adipocytes) of humans could be beneficial from a metabolic standpoint in settings of insulin resistance.

## AKNOWLEDGMENTS

We thank members of the Farese & Walther laboratory for helpful comments, and G. Howard for editorial assistance. Longwood small animal imaging facility at Beth Israel Deaconess Medical Center for PET/CT analysis. This work was supported by R01DK124913. T.C.W is an investigator of the Howard Hughes Medical Institute.

## AUTHOR CONTRIBUTIONS

C.C., R.V.F., and T.C.W. planned the study and designed the experiments. C.C. generated DGAT double flox and BA-DGAT KO mice. C.C. performed most of the experiments. A.F. analyzed mitochondrial respiration and, performed western blots of isolated mitochondria. C.C., R.V.F., and T.C.W. wrote the manuscript.

## DECLARATION OF INTERESTS

The authors declare no competing interests.

## STAR METHODS

- KEY RESOURSES TABLE
- CONTACT FOR REAGENT AND RESOURSE SHARING
- EXPERIMENTAL MODEL AND SUBJECT DETAILS
  - Generation of BA-DGAT KO mice
  - Mouse Husbandry
  - Cold exposure Studies
- METHOD DETAILS
  - Metabolic Tracer Studies
  - ^18^-FDG-PET/CT Analysis
  - RNA extraction and Quantitative Real-Time PCR
  - Immunoblotting
  - Comprehensive Lab Animal Monitoring System (CLAMS)
  - Tissue Lipid Extraction and Measurements
  - Mitochondrial Isolation
  - Seahorse Analysis
- QUANTIFICAION AND STATISTICAL ANALYSIS

## CONTACT FOR REAGENT AND RESOURCES SHARING

Further information and request for reagents and resources should be mailed to Robert V. Farese, Jr. and Tobias C. Walther.

## EXPERIMENTAL PROCEDURES

### Generation of BA-DGAT KO Mice

To generate BAT-specific *Dgat1* and *Dgat2* double knockout (BA-DGAT KO) mice, we first generated *Dgat1* and *Dgat2* double-floxed mice (D1D2 flox) by crossing *Dgat1*^*flox/flox*^ mice (Shih et al., 2009) (The Jackson Laboratory stock number: 017322) with *Dgat2*^*flox/flox*^ mice (Chitraju et al., 2019) (The Jackson Laboratory stock number: 033518). To generate BA-DGAT KO mice, we next crossed D1D2 flox mice with transgenic mice expressing Cre recombinase under control of mouse *Ucp1* promoter (Kong et al., 2014) (The Jackson Laboratory stock number: 024670).

### Mouse Husbandry

All mouse experiments were performed under the guidelines from Harvard Center for Comparative Medicine. Mice were maintained in a barrier facility, at room temperatures (22–23°C), on a regular 12-h light and 12-h dark cycle and had *ad libitum* access to food and water unless otherwise stated. Mice were fed on standard laboratory chow diet (PicoLab^®^ Rodent Diet 20, 5053; less than 4.5% crude fat) or Western-type high-fat diet (Envigo, TD.88137; 21.2% fat by weight, 42% kcal from fat).

### Cold Exposure Studies

For cold exposure experiments, mice were single-housed in the morning around 8:00 am. Mice had free access to food and water unless otherwise stated. Hourly core body temperatures were recorded using a rectal probe thermometer.

### Metabolic Tracer Studies

Mice were housed in cages without food for 1 hour before the start of the experiment. Mice were injected β-3 adrenergic receptor agonist CL 316,243 (intraperitoneal injection, 1 mg/kg body weight) or saline. After 1 hour of CL 316,243 injection, mice were intraperitonially-injected with [^3^H]-deoxyglucose (2 μCi/g bodyweight) or [^14^C]-oleic acid (0.5 μCi/g bodyweight) conjugated with bovine serum albumin. At 1 hour after tracer injection, mice were sacrificed by decapitation, and tissues were collected. Tissues were lysed in lysis buffer (0.25 M sucrose, 50 mM Tris HCl, pH 7.0, with protease inhibitor cocktail). Radioactivity in tissue lysates was measures by liquid scintillation counting.

### ^18^FDG-PET/CT Analysis

PET/CT imaging studies were performed on a Siemens Inveon PET/CT Multimodality System (Hao et al., 2013). In brief, mice were fasted for 1 hour before the experiment. Mice were injected β-3 adrenergic receptor agonist CL 316,243 (intraperitoneal injection, 1 mg/kg body weight). At 1 hour after CL 316,243 injection, an intravenous injection of ^18^F-FDG was made into the tail. Subsequently, the mouse was placed onto the imaging bed under 2% isofluorane anesthesia for the duration of imaging. After acquiring CT images at 80 kV and 500 mA with a focal spot of 58 mm, with a binning factor of 1:x, a whole-body PET scan was acquired. Co-registration of the reconstructed CT and PET images and image analysis were done using the manufacturer’s software. For PET quantification, the regions of interest (ROI) were selected using CT images as guides.

### RNA Extraction and Quantitative Real-Time PCR (qRT-PCR)

Total RNA from brown and white adipose tissues was isolated by Qiazol lysis reagent and using the protocol of the RNeasy Kit (Qiagen). Complementary DNA was synthesized using the iScript cDNA Synthesis Kit (Bio-Rad), and qPCRs were performed using the SYBR Green PCR Master Mix Kit (Applied Biosystems).

### Immunoblotting

Tissues were lysed using RIPA lysis buffer (25 mM Tris•HCl pH 7.6, 150 mM NaCl, 1% NP-40, 1% sodium deoxycholate, 0.1% SDS) containing protease inhibitors. Proteins were denatured in Laemmli buffer and were separated on 10% SDS-PAGE gels, and transferred to PVDF membranes (Bio-Rad). The membranes were blocked with blocking buffer for 1 hour in TBST containing 5% BSA or 5% milk, and then incubated with primary antibodies overnight. The membranes were then washed with TBST for 10 min x 3 times, and incubated in mouse secondary antibodies (Santa Cruz Biotechnology) at 1:5000 dilutions in blocking buffer. Membranes was washed again with TBST for 10 min x 3 times and revealed using the Super Signal West Pico kit (Thermo Scientific).

### Tissue Lipid Analysis

Brown adipose tissue was homogenized in 1 ml of lysis buffer (0.25 M sucrose, 50 mM Tris HCl, pH 7.0 with protease inhibitor cocktail). The homogenate was mixed with 5 ml of chloroform:methanol (3:2 v/v) and extracted for 2 hours by vigorous shaking. Upon centrifugation at 3000 x g at room temperature for 10 min, 100 µl of lower organic phase was collected and dried in a speed vac. To the dried lipids, 100– 300 µl of 0.1 % Triton X-100 was added and sonicated. Total TG content were measured using Infinity TM triglycerides reagent (Thermo Scientific) according to the manufacturer’s protocol. For plasma lipid measurements, 5 µl of plasma were used.

### Isolation of Mitochondria from Brown Adipose Tissue

Mitochondria were isolated from brown adipose tissue as described (Fischer et al., 2017; Luijten et al., 2019; Shabalina et al., 2013). Briefly, iBAT and scBAT of 2 mice were pooled in ice-cold isolation buffer (0.25 M sucrose buffer, 5 mM Tris pH 7.4, 2 mM EGTA, 2 % BSA). Tissues were minced, further homogenized in a glass homogenizer with 6 strokes, filtered through cotton gauze and centrifuged at 8500 g (JA-2550, Beckmann) for 10 min. The supernatant was discarded by inverting the tube, the pellet was resuspended in isolation buffer, homogenized in a glass homogenizer and centrifuged at 800 x g for 10 min. The supernatant, containing mitochondria, was centrifuged at 8500 x g for 10 min, and the mitochondrial pellet was resuspended in TES buffer (100 mM KCl, 20 mM TES, 1 mM EGTA, 0.6 % BSA, pH 7.2) to induce mitochondrial swelling. After centrifugation at 8500 g for 10 min, the supernatant was discarded and the mitochondria were resuspended in the remaining TES buffer. The solution was transferred to a small glass homogenizer, homogenized and the protein concentration was determined.

### Respiration Analysis

Respiration analysis was performed in a Seahorse XF96 analyzer (Agilent), as described before (Bartelt et al., 2018) with some modifications. A total of 5 μg mitochondrial protein in 20 μl TES buffer per well was loaded into Agilent Seahorse XF96 cell culture microplates and mitochondria were pelleted by centrifugation at 4000g for 15 min. Respiration was measured in respiration media (100 mM KCl, 20 mM TES, 1 mM EGTA, 2 mM MgCl_2_, 1 mM KH_2_PO_4_, 0.5 mM CaCl_2_, 0.5 % BSA, pH 7.2) containing substrate (20 mM glycerol-3-phosphate, 10 mM succinate + 5 μM rotenone or 5 mM pyruvate + 3 mM malate). UCP1-dependent respiration was blocked by addition of 5 mM GDP (Port A), and UCP1-dependent respiration was calculated by subtracting the respiration values after GDP-addition from basal uncoupled respiration in the presence of substrate. ATP-production was stimulated by addition of 1 mM ADP (Port B) and inhibited by addition of 1 μM oligomycin (Port C). ATP-synthesis activity was calculated by subtracting the respiration levels after oligomycin addition from the respiration levels after ADP addition. Mitochondrial respiration was then inhibited by addition of 1 μM antimycin A and 1 μM rotenone (Port D). The results from five independent mitochondrial preparations per genotype (mice housed at room temperature) were used for analysis with at least 8 technical replicates per isolation and condition. For the calculation of the total mitochondrial yield, the protein concentration in the mitochondrial preparations was multiplied with the total volume. Western blot analysis, 20 μg of protein were separated on 4-15% SDS-PAGE, transferred onto PVDF-membranes in a wet blotting system.

### Comprehensive Lab Animal Monitoring System (CLAMS)

Mice were housed individually, and acclimatized for two days. Oxygen consumption, carbon dioxide release, energy expenditure, and activity were measured using a Columbus Instruments’ Oxymax Comprehensive Lab Animal Monitoring System (CLAMS) system according to guidelines for measuring energy metabolism in mice (Tschop et al., 2011).

### Statistical Analyses

Data are presented as mean ± SD (standard deviation). Statistical significance was evaluated by unpaired two-tailed Student’s t-test or two-way ANOVA with repeated measures.

## Supplemental Information

**Figure S1.**
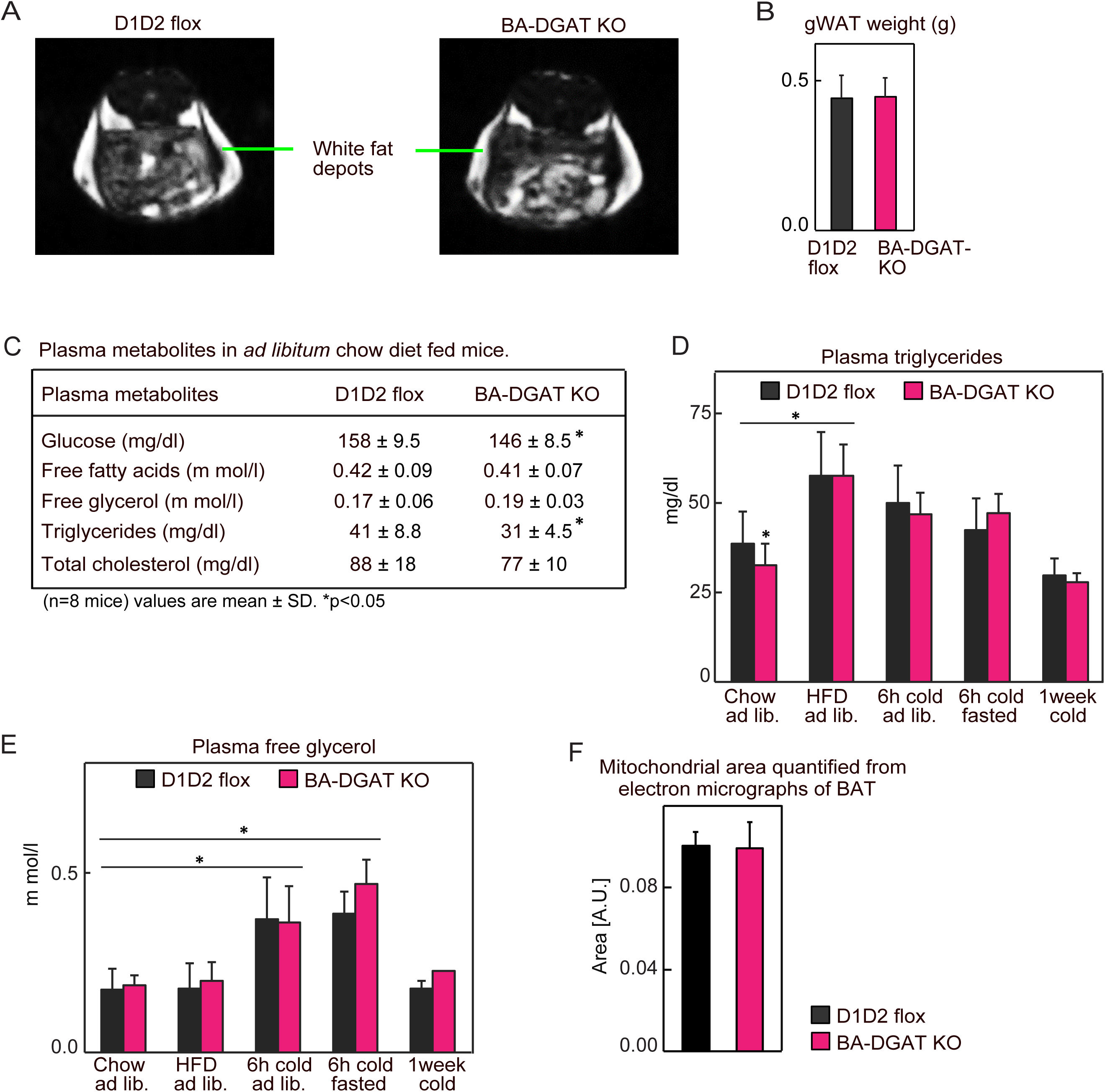
Related to Figure 1. BA-DGAT KO mice have normal white fat depots. (A) MRI scans of D1D2 flox and BA-DGAT KO mice (n=3 mice). (B) Weights of gonadal fat depots from chow-diet-fed mice (n=6). (C) Plasma metabolites in *ad libitum* chow-diet-fed mice (n=8 mice). (D) Plasma triglycerides (n=8 mice). (E) Plasma free glycerol (n=8 mice). (F) Mitochondrial area quantified from electron micrographs of BAT. Twenty mitochondria from three different sections of each genotype were quantified. Data are presented as mean ± SD. *p<0.05.

**Figure S2.**
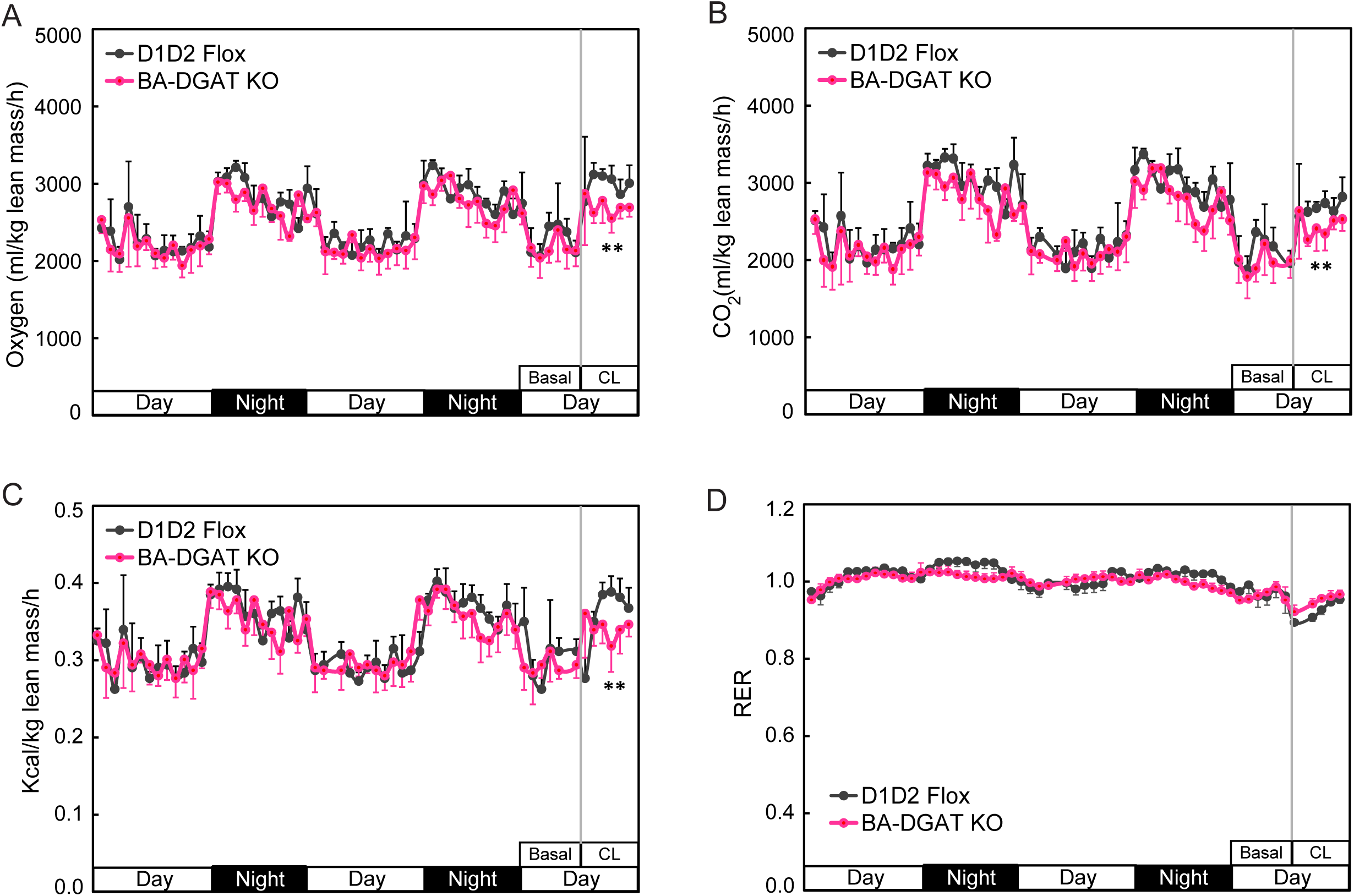
Related to Figure 2. Oxygen consumption, carbon dioxide release, energy expenditure and RER in basal and CL316243 injected mice. Each data point is mean of readings per hour (n=4 mice). Data are presented as mean ± SD. **p<0.01.

**Figure S3.**
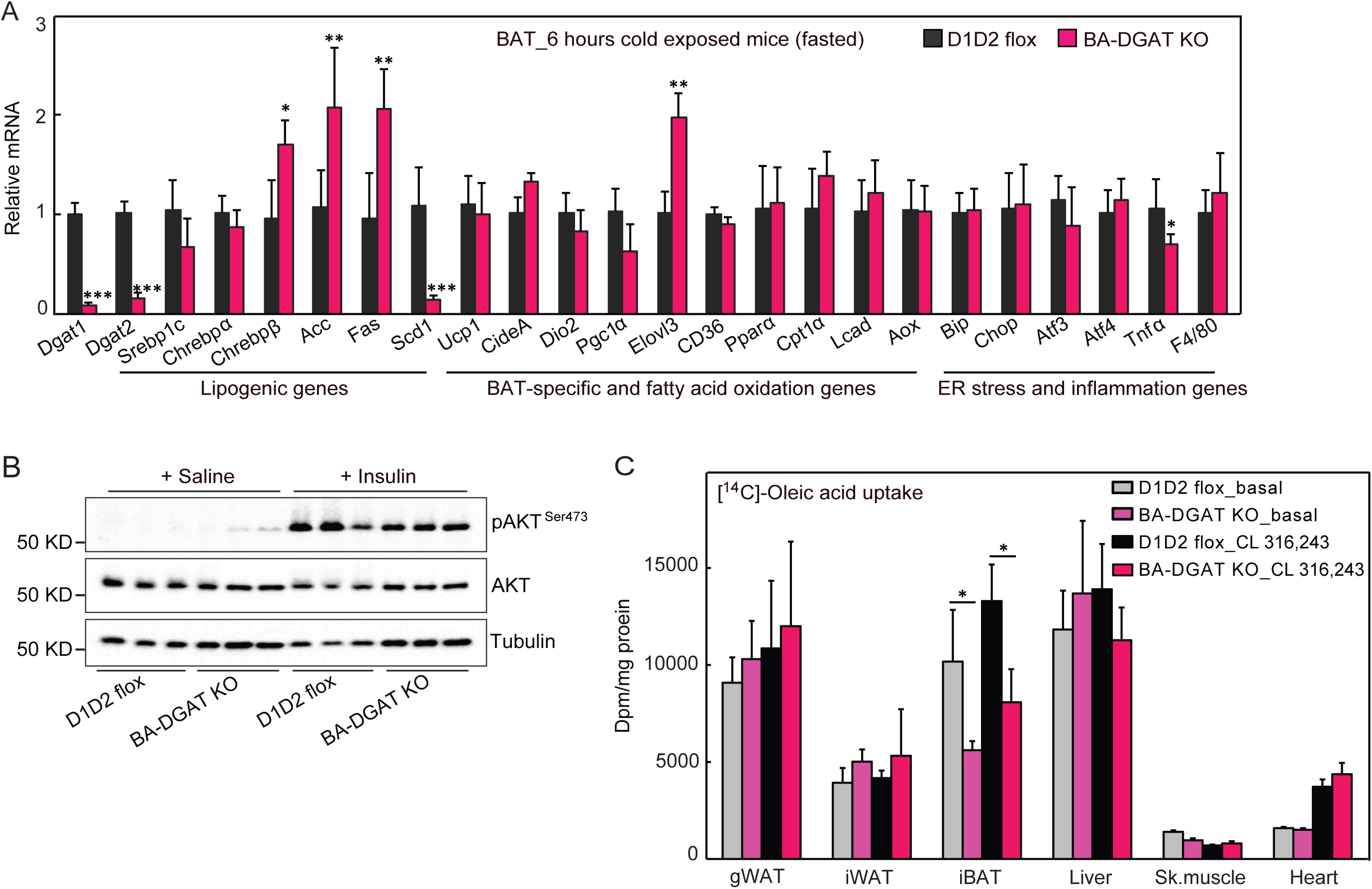
Related to Figures 2 and 3. (A) mRNA levels in BAT of cold-exposed (fasted) mice (n=6 mice). (B) Western blot analysis of insulin signaling in BAT. (C) [^14^C]-Oleic acid uptake by tissues in basal or in CL316243-injected mice (n=3 mice). Data are presented as mean ± SD. *p<0.05, **p<0.01, ***p<0.001.

**Figure S4.**
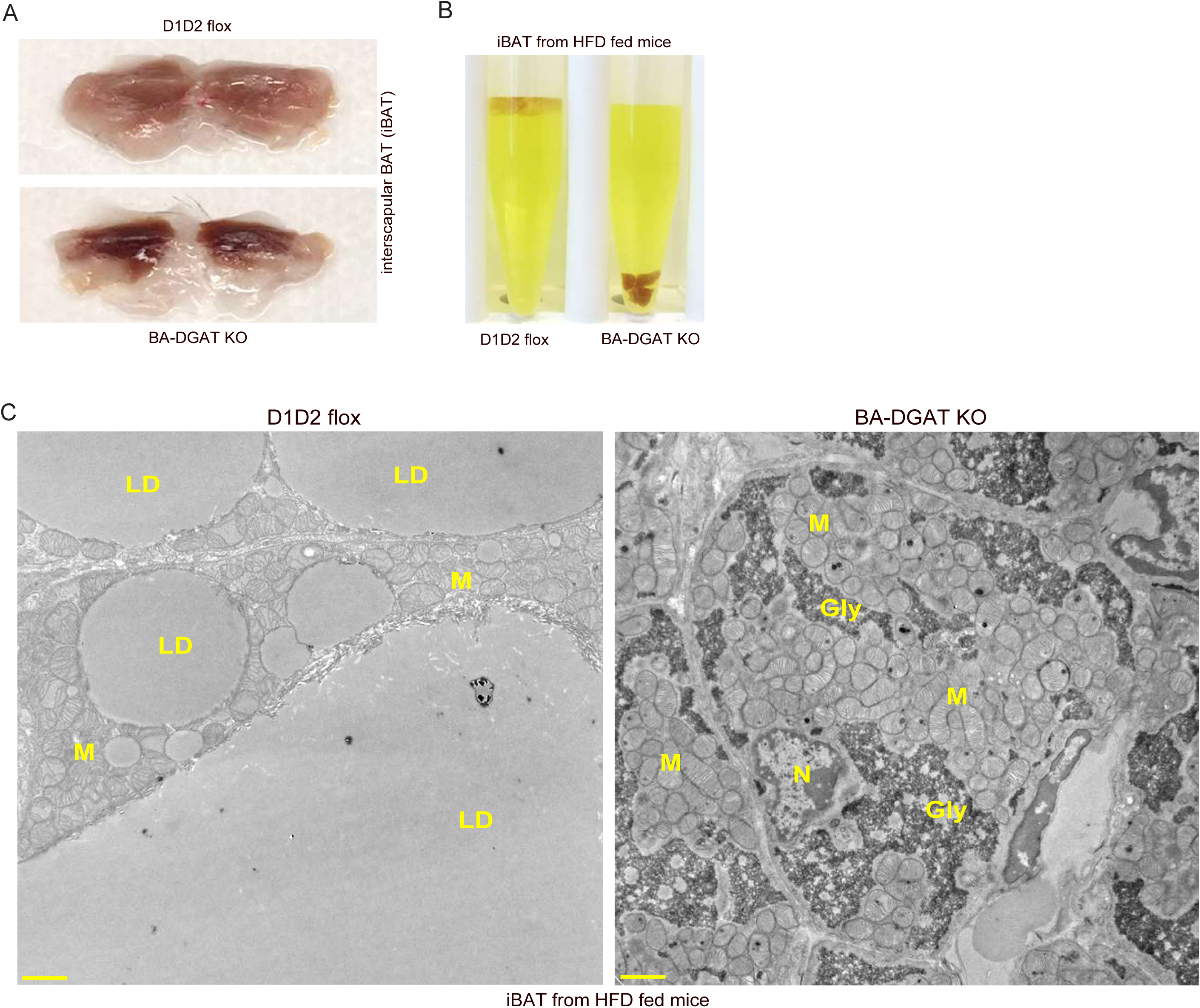
Related to Figure 4. (A) Gross appearance of iBAT from 12 weeks HFD-fed mice. (B) iBAT of HFD-fed BA-DGAT KO mice sinks in liquid fixative. (C) Brown adipose tissue of HFD-fed BA-DGAT KO mice stores glycogen (Gly). TEM images of iBAT from 12 weeks HFD-fed mice; scale bars, 2 μm. M, mitochondria; N, nucleus; Gly, glycogen.

